# Plant Cellulose as a Substrate for 3D Neural Stem Cell Culture

**DOI:** 10.1101/2023.06.30.547242

**Authors:** Lauren J. Couvrette, Krystal L. A. Walker, Tuan V. Bui, Andrew E. Pelling

## Abstract

Neural stem cell (NSC) based therapies are at the forefront of regenerative medicine strategies for various neural defects and injuries such as stroke, traumatic brain injury and spinal cord injury. For several clinical applications, NSC therapies require biocompatible scaffolds to support cell survival and to direct differentiation. Here, we investigate decellularized plant tissue as a novel scaffold for three-dimensional (3D) *in vitro* culture of NSCs. Plant cellulose scaffolds were shown to support attachment and proliferation of adult rat hippocampal neural stem cells (NSCs). Further, NSCs differentiated on the cellulose scaffold had significant increases in their expression of neuron-specific beta-III tubulin and glial fibrillary acidic protein compared to 2D culture on a polystyrene plate, indicating that the scaffold may enhance differentiation of NSCs towards astrocytic and neuronal lineages. Our findings suggest that plant-derived cellulose scaffolds have the potential to be used in neural tissue engineering and can be harnessed to direct differentiation of NSCs.

## 1. Introduction

Neural stem cells (NSCs) are self-renewing cells that proliferate *in vitro* and maintain the capacity to differentiate into neurons, astrocytes, and oligodendrocytes^1^. NSC transplantation is being investigated as a therapeutic strategy for numerous disorders of the central nervous system^2,3^ and many preclinical studies report promising results^4,5^. However, there are currently important limitations to the efficacy of NSC therapies such as low transplant survival and poor efficiency of neuronal differentiation. To overcome these issues, biocompatible scaffolds have emerged as a vehicle to engraft NSCs while supporting survival of the transplanted cells^6^. Cell scaffolds can enhance therapeutic efficacy of NSCs by promoting strong cell adhesion, guiding cell migration, and shielding transplanted NSCs from the cytotoxic injury environment occurring in CNS injuries^6,7^. Moreover, since NSC differentiation is mechanosensitive^8,9^, scaffolds’ physical properties such as porosity and elastic modulus can be exploited to influence stem cell differentiation^10–12^. It has been shown that stiffer matrices tend to be myogenic whereas softer matrices favor neuronal differentiation^13^ and promote expression of the neuronal marker βIII-tubulin in adult neural stem-cells^14^. Scaffold alignment also guides stem cell differentiation^15,16^ and studies indicate that anisotropic topographies support further axonal extension relative to isotropic topographies^17^.

In addition to physical cues, manipulating surface chemistry has been extensively investigated as a method to direct migration, proliferation, and differentiation of NSCs^18–21^. For instance, *Ge et al*. studied the effects of Poly-L-ornithine (PLO) on cell behavior *in vitro* and determined that NSCs cultured on PLO preferentially differentiated into neurons^22^ and had enhanced cell migration^23^. Likewise, NH_2_-terminated surfaces have been shown to promote neuronal differentiation of NSCs^20^. Further, some researchers have harnessed both topography and surface chemistry to direct stem cell behavior on scaffolds^24,25^.

In recent years, various three-dimensional cell culturing methods have been developed using scaffold-based technologies. These systems more accurately represent the *in vivo* microenvironment compared to traditional two-dimensional culture on polystyrene^26^ and the behavior of cells within 3D systems is more physiologically relevant, making it a better cell culturing method for tissue engineering and drug discovery^27–32^. As such, various scaffold-based neural constructs have been developed for use in disease modeling, regenerative medicine, and the study of the stem cell niche^33–36^.

Various researchers have investigated the possibility of producing plant-derived cellulose biomaterials, which were shown to support cell infiltration and vascularization *in vivo*^37–39^. In recent years, a range of plant tissues have been decellularized to produce biocompatible scaffolds for culturing mammalian cells^40–42^ as well as pre-clinical applications including skin tissue^43–45^, neural tissue^46^, and bone tissue engineering^47–50^. For example, spinach leaves were decellularized while maintaining their vascular architecture, which was then repopulated with human dermal microvascular endothelial cells^45^. Furthermore, *Flammulina velutipes* mushroom has been successfully used as a nerve guidance conduit in a rat model of sciatic nerve defect^46^. As evidenced above, plant-derived biomaterials are becoming increasingly attractive for biomedical applications, which can be attributed in part to improved cost effectiveness, scalability, and lower immunogenicity relative to animal sources.

Here, we investigate the viability of a plant-derived biomaterial as a 3D *in vitro* culture system for adult rat neural stem cells. We hypothesized that the stalks of *Asparagus officinalis*, which are composed of linearly arranged microchannels, would produce a scaffold with an interesting surface topology for the culture of neural stem cells, as topology is known to influence NSC differentiation and axon guidance. We first examined the asparagus scaffold’s physical characteristics by scanning electron microscopy and mechanical testing. Next, the scaffold’s ability to support attachment and migration of NSCs was assessed for various time periods. Further, we examined the differentiation potential of neural stem cells in this 3D culture system by immunostaining for markers including neuron-specific β-III tubulin and glial fibrillary acidic protein.

## 2. Methods

### 2.1 Biomaterial Production

Asparagus scaffolds were prepared utilizing decellularization methods described previously^51,52^. Using a 4 mm biopsy punch, Asparagus officinalis sections were cut and placed into a 50ml Falcon tube containing 0.1% sodium dodecyl sulphate (SDS) (Sigma-Aldrich). Samples were shaken for 72 hours at 180 RPM at room temperature. The resulting cellulose scaffolds were then transferred into new sterile microcentrifuge tubes, washed and incubated for 12 hours in phosphate-buffered saline (PBS). Following the PBS washing steps, the asparagus were then incubated in 100 mM CaCl2 for 24 hours at room temperature and washed 3 times with dH2O. Samples were then sterilized in 70% ethanol overnight. Finally, they were then washed 12 times with sterile 1X PBS.

### 2.2 Scanning Electron Microscopy

Scanning electron microscopy was performed at the 2-week timepoint. NSC-seeded scaffolds were fixed with 4% PFA and dehydrated through successive gradients of ethanol (50%, 70%, 95% and 100%). Samples were dried with a critical point dryer (SAMDRI-PVT-3D) then gold-coated at a current of 15mA for 3 minutes with a Hitachi E-1010 ion sputter device. SEM imaging was conducted at voltages ranging from 2.00–10.0 kV on a JSM-7500F Field Emission SEM (JEOL).

### 2.4 Mechanical Testing

After 7 days in culture media at 37°C, scaffolds (4mm diameter x 1.2mm height) were placed onto a CellScale UniVert (CellScale) compression platform for tensile testing. Each scaffold (n=15) was compressed mechanically to a maximum 30% strain, at a compression speed of 50 μm/s. The elastic modulus was determined from the slope of the linear region of the resulting stress-strain curves.

### 2.5 Cell culture, scaffold seeding and differentiation

The resulting cellulose scaffolds were incubated in poly-L-ornithine (Sigma, 20μg/ml in dH2O) overnight at room temperature. PLO-coated scaffolds were rinsed twice with sterile water before being transferred into a 96-well plate. Rat adult hippocampal neural stem cells (Sigma SCR022) were cultured in serum free medium (1X KnockOut D-MEM/F-12 with 2% StemPro Neural Supplement (ThermoFisher A1050801), 20 ng/mL bFGF, 20 ng/mL EGF, and 2mM GlutaMAX-I). Culture vessels were also coated with 20 μg/ml poly-L-ornithine and 10 μg/ml laminin as described above. An 80 μL droplet containing 200 000 cells was deposited onto every scaffold, which was then incubated at 37°C and 5% CO^2^. After 4 hours of incubation, 2 mL of StemPro NSC SFM complete medium was added to each scaffold, which was then incubated for 48 hours before transferring scaffolds to a new 96-well plate. For 2-4 weeks, media was exchanged daily. For neuronal and astrocyte differentiation, cells were cultured in 1X KnockOut DMEM/F-12 supplemented with 2% B27 (cat. 17504044 ThermoFisher), 2mM GlutaMAX-I (ThermoFisher)) for 7 days.

### 2.5 Staining and Confocal Microscopy

Cell attachment and morphology was documented by phase contrast microscopy at day 3, 7, and 14 post seeding. Staining and confocal microscopy was performed at 72h or 14 days in culture. NSC-seeded scaffolds were fixed with warm 4% paraformaldehyde for 10 minutes then incubated 3 minutes in warm permeabilization buffer (0.5% Triton-X, 20 mM HEPES, 300mM Sucrose, 50mM NaCl, 3mM MgCl2, 0.05% sodium azide). Samples were then incubated for 15 minutes in Fluorescein Phalloidin (1:100, ThermoFisher F432) to stain F-actin. Samples were rinsed with PBS and incubated in Hoechst (1:200, ThermoFisher) for 10 minutes to label nuclei. Scaffolds were incubated in 0.2% Congo Red (Sigma) for 15 minutes before a final PBS rinse.

For immunostaining GFAP and β-tubulin, cells or cell-seeded scaffolds were fixed in 4% paraformaldehyde for 15 minutes at room temperature. Samples were incubated for 5 minutes in permeabilization buffer (0.5% Triton-X, 20 mM HEPES, 300mM Sucrose, 50mM NaCl, 3mM MgCl2, 0.05% sodium azide). After blocking in 6% Normal Goat Serum in 1X PBS for 10 minutes, samples were incubated in rabbit anti-GFAP antibody (cat. AB5804 Sigma, 1:1000 in 1X PBS) or mouse anti-β-III tubulin antibody (cat. MAB1195 R&D systems, 10ug/ml in 1X PBS) overnight at 4°C. The following day, samples were washed twice in 1xPBS (5 mins, RT) followed by a 2-hour incubation in secondary antibody: goat anti-rabbit IgG Alexa Fluor 488 (cat. A11008 Invitrogen, 1:500 in 1X PBS) or goat anti-mouse IgG Alexa Fluor 594 (cat. A11005 Invitrogen, 1:200 in 1X PBS). Samples were then washed in 1X PBS and counterstained with Hoescht 33342 (1:2000 in 1X PBS) for 10 minutes All samples were then mounted in Vectashield (Vector Labs) and imaged on with a Nikon A1R laser scanning confocal microscope with appropriate filter sets and laser lines.

### 2.6 Image analysis

For ßIII-tubulin stains, analysis was performed on 13 images at 40X magnification for each condition (2D and 3D). Total cell number was determined by cell counting Hoescht-labeled nuclei using FIJI. Cells were considered ßIII-tubulin^+^ when signal from red and blue channels were colocalized. For GFAP stains, analysis was performed on 6 images at 40X magnification for each condition (2D and 3D). Cells were considered GFAP^+^ when signal from green and blue channels were colocalized.

### 2.7 Alamar Blue Cell Proliferation Assay

To measure cell proliferation, an Alamar blue assay was performed according to the manufacturer’s protocol. Briefly, PLO-coated asparagus scaffolds were placed into the wells of a 96-well plate and seeded with 100 000 adult rat hippocampal neural stem cells. After 1, 2 or 5 days, cell-seeded scaffolds were transferred into a new well and 200μL of fresh media containing 10% Alamar blue (cat. BUF012A Bio-Rad) was added to each scaffold. After 4 hours of incubation at 37°C, 100μL of media was removed from each well and deposited into an empty well before reading absorbance at 570nm and 600nm on a spectrophotometer (Epoch 2, BioTek). Each timepoint included biological triplicates for cell-seeded scaffolds (n=3), 2D controls (NSCs grown on a PLO-coated well, n=3) and blanks (media only, n=3). Absorbances were corrected by subtracting the average absorbance of blanks at 570nm & 600nm.

### 2.8 Statistical Analysis

P values were calculated using two-tailed Student’s t test. Results were considered as statistically significant when P < 0.01. Numerical data are expressed as mean ± standard deviation.

## 3. Results

### 3.1 Characterization of plant cellulose scaffold

Raw *Asparagus officinalis* stalks were cut into discs of 4mm in diameter and 1.2mm in height, which were then decellularized to remove all native cells. The resulting scaffold is composed of vascular bundles (VBs) interspersed between parenchyma (Fig. 1A). Naturally occurring structures within the scaffold were characterized by scanning electron microscopy (Fig. 1B-D). A majority of the scaffold’s surface consists of parenchyma that have an average pore diameter of 39±15μm. In addition, each scaffold was found to contain 14±2 vascular bundles, which are aligned channels that traverse the length of the scaffold, with an average spacing of 602±61μm. The distribution of channel diameters observed in these scaffolds may be conducive to cell survival, since such porous networks have previously been demonstrated to enable nutrient exchange and waste removal within scaffolds^53,54^. The Young’s Modulus of the cellulose scaffold in culture media at 37°C is 128±20 kPa (n=5) when measured parallel to the long axis. Moreover, the scaffolds were coated overnight in poly-L-ornithine (PLO) to promote cell attachment. Surface modification with PLO was confirmed with SEM (Fig. 1D).

**Figure 1.**
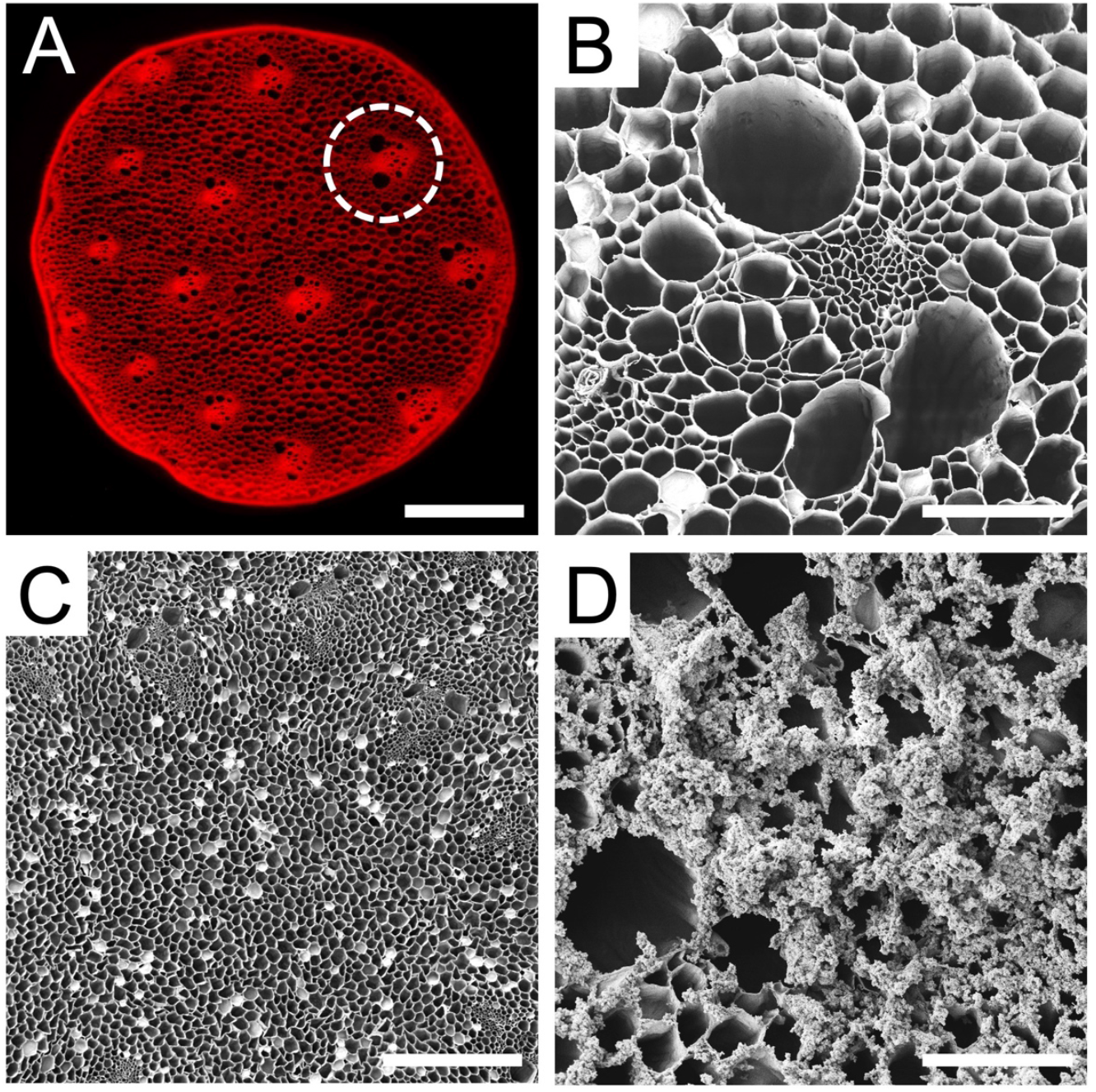
Cross sections of decellularized *Asparagus officinalis* scaffolds. **A**. Confocal microscope maximum intensity projection of a decellularized asparagus scaffold stained with Congo Red for cellulose. The decellularization process preserves vascular bundles (VBs, circled), which are microchannels that run along the asparagus stalk and separated by porous parenchyma tissue (scale bar = 1mm) **B**. High magnification SEM of an individual VB within scaffold which reveals the long microchannels (scale bar = 100μm) **C**. SEM of the parenchyma tissue which reveals numerous pores with a wide distribution of diameters (scale bar = 500μm) **D**. Higher magnification SEM of a scaffold coated with poly-L-ornithine, a positively charged synthetic amino acid which gives the scaffold surface a sabulous appearance (scale bar = 100μm). Such appearance is never observed on uncoated decellularized scaffolds and only appears after PLO coating.

### 3.2 Cellulose scaffold supports rat NSC attachment and proliferation

A single-cell suspension of NSCs was seeded onto PLO coated scaffolds and cultured for 3 to 14 days. As early as 3 days in culture, neurospheres had attached to the cellulose scaffold and were visualized through F-actin staining (Fig. 2A). Many neurospheres with diameters of 50μm to 300μm were found on the surface of the scaffold (Fig. 2B). By 14 days in culture the same high density of neurospheres on the scaffolds was observed to persist (Fig. 2C). In order to examine how the NSCs penetrated into the VB microchannels, the scaffold was sectioned longitudinally along it long axis. This allowed us to visualize the migration of NSCs and neurospheres down the cellulose channels (Fig. 2D). Importantly, NSC cells and neurospheres were observed inside of the scaffold as well as on its surface. Finally, we also observed that NSCs were able to migrate out of the attached neurospheres (Fig. 2E). Groups of cells were observed migrating and extending out from several neurospheres onto the scaffold. In addition to microscopic examination, cell proliferation was also assessed. This was achieved on the scaffolds with an AlamarBlue assay, which monitors the chemical reduction of culture media to detect metabolic activity of cells. Over 5 days in culture (Fig. 3), the percentage of reduced AlamarBlue reagent increased progressively, indicating continued growth of NSCs on the scaffold. When compared to NSCs grown as a monolayer on a polystyrene culture plate, cells on the scaffold had a slight reduction in metabolic activity during the first 5 days of growth but exhibited a similar trend.

**Figure 2.**
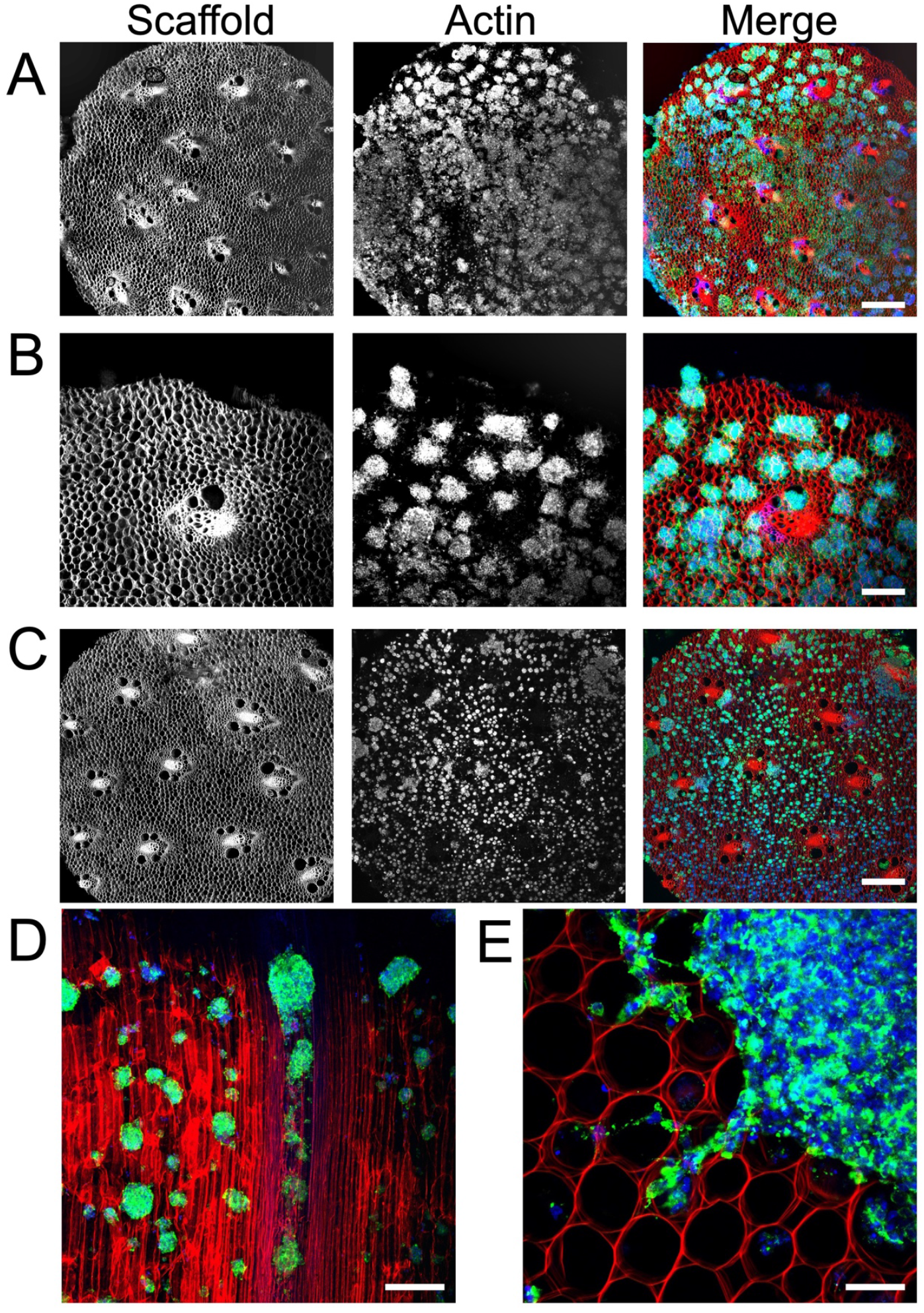
Confocal microscopy (maximum intensity projection) of adult rat NSCs cultured on decellularized, PLO coated asparagus scaffold. F-actin was stained with phalloidin (green), nuclei were stained with Hoechst 33342 (blue) and the cellulose scaffold was stained with congo red (red). **A**. 4X magnification of NSCs grown on scaffold for 3 days revealing numerous neurospheres (scale bar = 500μm). **B**. 10X magnification cross section of NSCs on scaffold at 3 days in culture shows the distribution of neurospheres at greater detail (scale bar = 100μm). **C**. 4X magnification of NSCs grown on scaffold for 14 days reveals the continued presence and distribution of neurospheres on the scaffold surface (scale bar = 500μm). **D**. After sectioning the scaffold longitudinally along its long axis (parallel to the direction of the VB microchannels) a 10X magnification image reveals the NSCs migrating into scaffold channels at 3 days in culture (scale bar = 200μm). The highly aligned structure of the cellulose scaffold is easily observed (red) and the presence of NSCs and neurospheres are observed deep within the scaffold. **E**. 40X magnification cross section of NSCs grown on scaffold for 14 days (scale bar = 50μm) reveals groups of NSCs migrating and projecting out of the edge of a single neurosphere.

**Figure 3.**
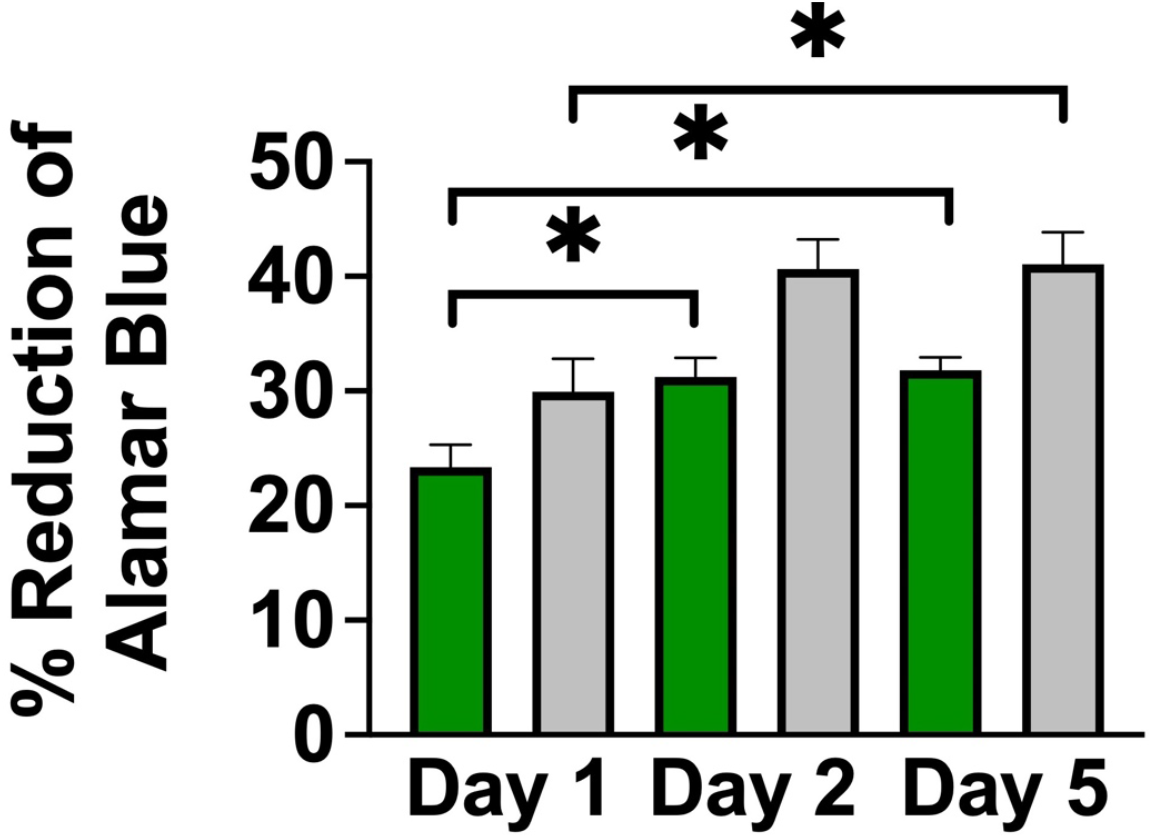
Evaluation of cell proliferation by Alamar Blue assay. The percent reduction of AlamarBlue reagent by NSCs grown on a cellulose scaffold (in green) compared to a polystyrene culture plate (in grey) over 5 days in culture. Metabolic activity of cells increased over 5 days in culture, indicating continued growth of NSCs on the scaffold. Statistical significance (* indicates p<0.01) was determined using a student’s t-test. (Error bars represent standard deviation, N=3 for each condition)

### 3.3 Plant cellulose scaffold enhances neuronal and astrocytic differentiation

To determine the effects of this 3D culture system on NSC differentiation potency, we monitored the expression of lineage specific markers. A single-cell suspension of NSCs was simultaneously seeded onto PLO-coated scaffolds and onto a PLO-coated culture plates at equal seeding densities. After seven days in differentiation media, cells were fixed and immunostained for glial fibrillary acidic protein (GFAP, an astrocytic marker) or neuron specific ßIII-tubulin (immature neuron marker). Immunostaining revealed a significantly higher fraction of GFAP positive cells were observed on the cellulose scaffold (18.45±2.8%) compared to the 2D monolayer on a polystyrene culture plate (3.50±2.7%) (Fig. 4A-C) (P<0.01, n=6). Similarly, immunostaining revealed enhanced expression of ßIII-tubulin on the 3D scaffold (Fig. 4D-F). Cells in 2D condition were 0.79±0.7% ßIII-tubulin positive whereas cells grown on the 3D scaffold were 16.46±4.5% ßIII-tubulin positive, which represents a significant increase in expression of this early neuronal marker (P<0.001, n = 13). Both findings demonstrate that NSCs retain their ability to differentiate into various lineages when cultured in this 3D cellulose scaffold.

**Figure 4.**
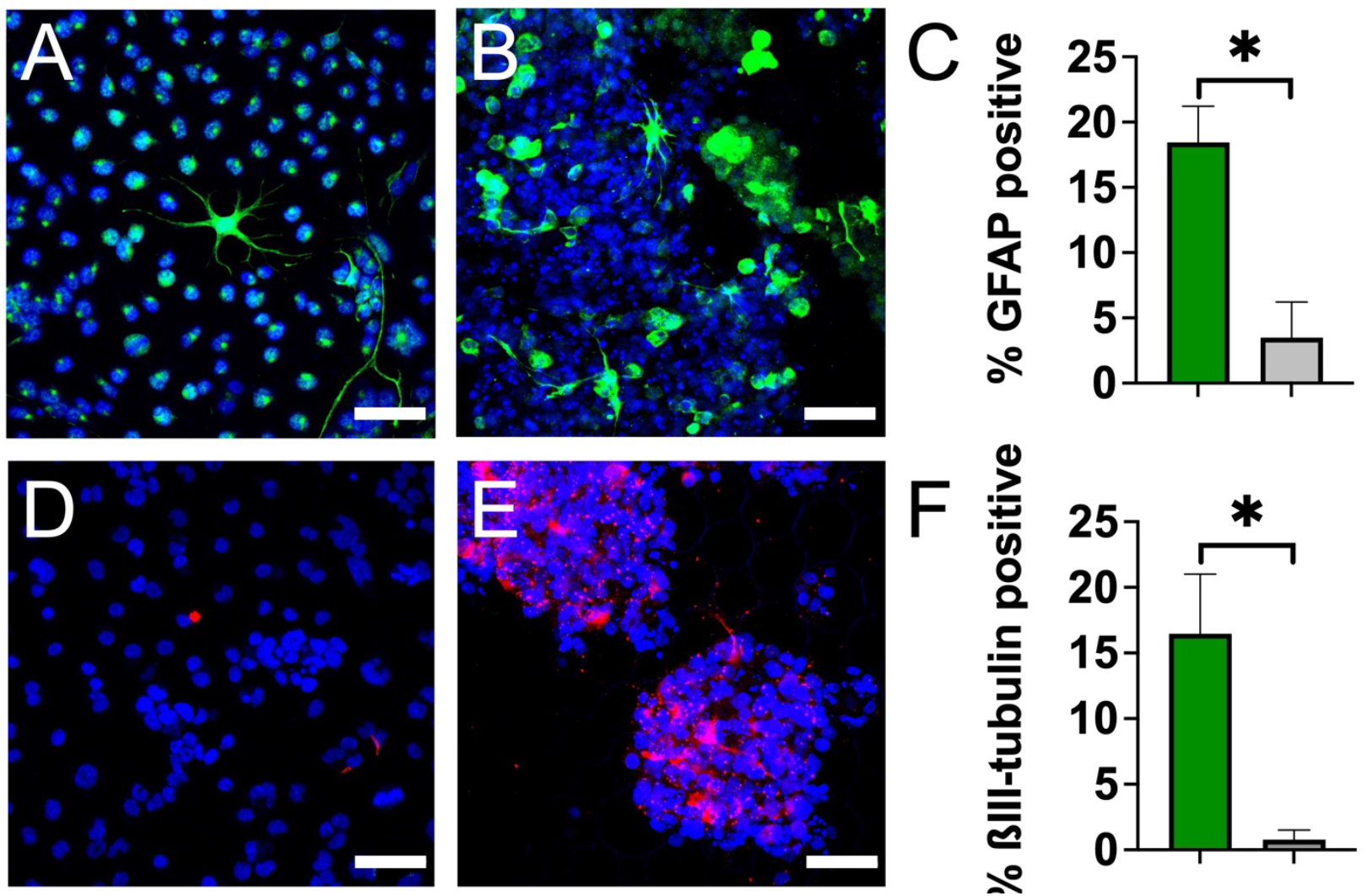
NSC lineage analysis by immunostaining reveals enhanced neuronal and astrocytic differentiation on cellulose scaffold. Representative confocal microscopy (maximum intensity projections) of adult rat NSCs after 7 days in culture in differentiation media. Nuclei were stained with Hoechst 33342 (blue). **A**. NSCs grown on PLO-coated culture plates (2D) stained for GFAP (green) (scale bar = 50μm). GFAP positive cells were identified as possessing green signal throughout the entire cell body and not just localized to the nucleus **B**. NSCs grown on PLO coated scaffold (3D) stained for GFAP (green) (scale bar = 50μm). **C**. Percentage of GFAP positive adult rat NSCs after 7 days in culture on 3D scaffolds (in green) compared to 2D polystyrene plates (in grey). Statistical significance (* indicates p<0.01) was determined using a student’s t-test. **D**. NSCs grown on PLO-coated culture plates (2D) stained for ßIII-tubulin (red) (scale bar = 50μm). **E**. NSCs grown on PLO-coated scaffold (3D) stained for ßIII-tubulin (red) (scale bar = 50μm). **F**. Percentage of ßIII-tubulin positive adult rat NSCs after 7 days in culture on 3D cellulose scaffold (in green) compared to 2D polystyrene plate (in grey). Statistical significance (* indicates p<0.01) was determined using a student’s t-test.

## 4. Discussion

In this study, we developed a 3D cell culture scaffold consisting of decellularized *Asparagus officinalis* stalks that supported attachment, proliferation, and differentiation of rat adult neural stem cells. After decellularization, the cellulose scaffold was seeded with primary neural stem cells isolated from the hippocampus of adult Fisher 344 rats. Microscopic examination of the scaffolds revealed a system of aligned channels with various diameters, which we predicted would allow for efficient transport of nutrients and provide guidance cues for cell attachment. Our results demonstrate that within 3 days, NSCs attached to the scaffold, both as individual cells and neurospheres of diverse sizes (Fig. 2). Groups of NSCs appear to migrate out from the neurospheres attached to PLO-coated scaffolds. This behavior can be explained by the fact that PLO enhances migration by promoting filopodia formation^23^. Subsequent F-actin staining revealed the morphology of neurospheres within the biomaterial and highlighted the migration of NSCs into the channels of the scaffold. Further, NSCs were found to proliferate within the 3D culture system, as demonstrated by an Alamar Blue assay (Fig. 3). Taken together, these data suggest the scaffold is biocompatible and has appropriate physical characteristics to allow for neural stem cell growth.

The behavior of cells cultured in 3D systems is often different from that in 2D, including the rate of proliferation^55,56^. Interestingly, the percent reduction in AlamarBlue reagent was consistently lower in the 3D scaffold compared to the 2D monolayer culture system, suggesting a slight inhibition of NSC growth on the scaffolds. This difference can be attributed in part to the increased heterogeneity in the 3D culture system, where NSCs assembled into neurospheres attached to the scaffold. While the outer surface of these neurospheres consists of cells with high rates of proliferation, the inner layers of NSCs tend to be quiescent or necrotic due to reduced access to oxygen, nutrients, and growth factors. Therefore, this difference in reduction of AlamarBlue may result from heterogeneity rather than purely from differences in the rate proliferation.

Finally, NSCs grown on the scaffolds were cultured in differentiation media and their expression of cytosolic markers GFAP & ßIII-tubulin was evaluated after 7 days. Remarkably, this plant-derived scaffold appears to enhance NSC differentiation towards astrocytes and neurons, as evidenced by significant increases in GFAP positive and ßIII-tubulin positive cells on the scaffold compared to 2D controls (Fig. 4). It is well established that the differentiation of stem cells can be influenced by several extrinsic and intrinsic factors including chemical cues, the physical characteristics of the environment, and the surrounding cell types^15–17^. Therefore, the differences observed in our differentiation experiments are likely due to a complex interplay of cues provided by the scaffold. An important factor in guiding the fate of NSCs is scaffold architecture and anisotropy. It was previously reported that anisotropic contact promotes neuronal differentiation of NSCs^57^. In accordance, we propose that the plant scaffold’s aligned channels provide geometric cues that may modulate differentiation toward higher neurogenesis. In addition, scaffold elasticity is another key factor which mediates stem cell fate^33^. To determine the elastic modulus of the scaffold, mechanical tests were performed after 1 week in media at 37°C. The Young’s modulus of the scaffold was determined to be 128±20 kPa, which is softer than polystyrene cell culture plates (E = 3.73 GPa)^58^ but slightly stiffer than brain and spinal cord tissue. Previous studies have shown that softer growth substrates tend to favor neuronal differentiation^13,14,59^. Here, the difference in stiffness between the scaffold and the polystyrene cell culture plate likely contributed to the increase in neuronal differentiation. As well, the 3D growth environment may have enhanced cell-cell signaling for lineage differentiation, which could induce more differentiation of NSCs on the scaffold relative to 2D culture, as others have reported^60–62^.

Interestingly, we observed many neurospheres forming in the confined circular pores of the parenchyma tissue (Fig 2A-C). In previous work, we have shown how embryonic stem cell spheroid formation can be stimulated by the physical confinement of cells in small microscale environments^63^. This is not dissimilar to what takes place in traditional hanging drop and round bottom multi-well culture models designed for stimulating spheroid formation^64^. In such environments, cells become trapped and interact more frequently which nucleates the formation of a spheroid. Here, we speculate that a similar phenomenon is taking place leading to a significant increase in the number of spheroids on and inside of the scaffold as opposed to on a flat substrate. However, at this point the precise mechanism which leads to enhanced NSC differentiation on these scaffolds is unclear and future work will be required to understand this phenomenon. In summary, we have demonstrated that cellulose scaffolds support the growth and differentiation of neural stem cells *in vitro*. Our findings suggest that plant derived scaffolds could facilitate the production of large numbers of specifically differentiated cells needed for NSC research or regenerative medicine.

## Author Contributions

L.J.C., K.L.A.W. and A.E.P. designed the study. L.J.C. and K.L.A.W. contributed to experimental design and execution; data acquisition, review, and analysis; and manuscript preparation and revision. All authors participated in writing the manuscript. All authors have read and agreed to the published version of the manuscript.

## Funding

This project was supported by grants from the Li Ka Shing Foundation and the Natural Sciences and Engineering Research Council of Canada.

## Institutional Review Board Statement

Not applicable

## Informed Consent Statement

Not applicable

## Data Availability Statement

The data presented in this study are available on request from the corresponding author.

## Conflicts of Interest

L.J.C., K.L.A.W and A.E.P. are inventors on a patent application which describes PLO coating of plant based biomaterials.

